# Systems biology-enabled targeting of NF-κΒ and BCL2 overcomes microenvironment-mediated BH3-mimetic resistance in DLBCL

**DOI:** 10.1101/2024.11.30.626166

**Authors:** Aimilia Vareli, Haripriya Vaidehi Narayanan, Heather Clark, Eleanor Jayawant, Hui Zhou, Yi Liu, Lauren Stott, Fabio Simoes, Alexander Hoffmann, Andrea Pepper, Chris Pepper, Simon Mitchell

**Affiliations:** Department of Clinical and Experimental Medicine, Brighton and Sussex Medical School, Falmer, United Kingdom; Department of Clinical and Experimental Medicine, Brighton and Sussex Medical School, Falmer, United Kingdom; Signaling Systems Laboratory, Department of Microbiology, Immunology, and Molecular Genetics, and Institute for Quantitative and Computational Biosciences, University of California, Los Angeles, Los Angeles, CA, United States; DeepKinase Biotechnologies, Beijing, China

## Abstract

In Diffuse Large B-cell Lymphoma (DLBCL), elevated anti-apoptotic BCL2-family proteins (e.g., MCL1, BCL2, BCLXL) and NF-κB subunits (RelA, RelB, cRel) confer poor prognosis. Heterogeneous expression, regulatory complexity, and redundancy offsetting the inhibition of individual proteins, complicate the assignment of targeted therapy. We combined flow cytometry ‘fingerprinting’, immunofluorescence imaging, and computational modeling to identify therapeutic vulnerabilities in DLBCL. The combined workflow predicted selective responses to BCL2 inhibition (venetoclax) and non-canonical NF-κB inhibition (Amgen16). Within the U2932 cell line we identified distinct resistance mechanisms to BCL2 inhibition in cellular sub-populations recapitulating intratumoral heterogeneity. Co-cultures with CD40L-expressing stromal cells, mimicking the tumor microenvironment (TME), induced resistance to BCL2 and BCLXL targeting BH3-mimetics via cell-type specific upregulation of BCLXL or MCL1. Computational models, validated experimentally, showed that basal NF-κB activation determined whether CD40 activation drove BH3-mimetic resistance through upregulation of RelB and BCLXL, or cRel and MCL1. High basal NF-κB activity could be overcome by inhibiting BTK to resensitize cells to BH3-mimetics in CD40L co-culture. Importantly, non-canonical NF-κB inhibition overcame heterogeneous compensatory BCL2 upregulation, restoring sensitivity to both BCL2- and BCLXL-targeting BH3-mimetics. Combined molecular fingerprinting and computational modelling provides a strategy for the precision use of BH3-mimetics and NF-κB inhibitors in DLBCL.

## Introduction

Diffuse large B-cell lymphoma (DLBCL), is an aggressive blood cancer (1). First line treatment is only successful in approximately 60% of patients (2–5). The highly heterogenous nature of the disease has challenged advancing the standard of care through one-size-fits-all approaches (6–11). Activated B-cell (ABC) DLBCL patients bearing Nuclear Factor-κappa light chain enhancer of Β cells (NF-κB) activating mutations, and Germinal Center (GC) DLBCL patients with *BCL2*-activating mutations have particularly poor prognosis (11).

NF-κΒ is a dimeric transcription factor composed of combinations of five proteins (RelA, RelB, cRel, p50 and p52), which are normally sequestered in the cytoplasm by a combination of four inhibitors of NF-κΒ (IκBα, -β, -δ, and-ɛ) (12, 13). The IκB proteins have distinct affinities for different NF-κB dimers with IκBα predominantly inhibiting RelA-containing dimers and IκB-δ and -ɛ regulating cRel-containing dimers in a context-specific manner (14, 15). The abundance and nuclear localization of individual NF-κΒ components is prognostically informative in DLBCL (16–18). Heterogeneity in NF-κB subunits can be quantified and visualized using flow cytometry to reveal unique ‘fingerprints’ of NF-κB, with heterogenous levels of individual NF-κB proteins found within ABC-DLBCL (19). Two NF-κΒ pathways (canonical and non-canonical) may be activated by a combination of tumor microenvironmental signaling and activating mutations in DLBCL leading to nuclear translocation of NF-κΒ (20). While targeting NF-κB has shown therapeutic potential in subgroups of blood cancer patients (21–26), heterogeneity and toxicities associated with global targeting of the canonical NF-κB pathway has challenged clinical progress. The oncogenic role of NF-κΒ in lymphoma is largely attributed to the anti-apoptotic gene expression induced upon NF-κΒ activation (26, 27).

Anti-apoptotic BCL2 family proteins, including Myeloid Cell Leukemia 1 (MCL1) and B-cell Lymphoma-extra Large (BCLXL), are inhibitors of the intrinsic pathway of apoptosis (28, 29), and are often overexpressed in DLBCL as a result of gene translocation events, gene amplifications, mutations within open reading frames, and microenvironment-mediated increased NF-κB activity (30). BCL2 is an attractive target in DLBCL, and the specific BCL2 inhibitor venetoclax is clinically used for multiple hematological malignancies (31). However, venetoclax has shown limited efficacy in DLBCL clinical trials (31, 32). BCL2-family proteins (MCL1 and BCLXL) are often over expressed in DLBCL with elevated expression conferring venetoclax resistance by compensating for the sequestration of BCL2 (33, 34). BH3-profiling, established in 2006, measured the impact of BH3-peptides on mitochondrial outer membrane permeabilization (35). While predictive of response to BCL2 antagonism in pre-clinical studies, responses to BH3-mimetics in drug response assays can differ, and BH3-profiling has not translated into prospective assignment of therapies in clinical trials of lymphoma (36–38).

The daunting complexity of both NF-κΒ regulatory networks, and apoptotic signaling regulation, has motivated the use of computational systems biology models to predict responses to BH3-mimetics (39), responses to the tumor microenvironment (TME) (19, 40), and predict the impact of mutations on clinical outcome (19, 41). Recent single-cell RNA sequencing of DLBCL patient material revealed the primary impact of the TME is through CD40 and B-cell activating factor (BAFF), which activate non-canonical NF-κB (25). The regulation of BCLXL by non-canonical NF-κB (RelB) is established in CLL (25). However, how heterogeneity in multiple NF-κB proteins results in anti-apoptotic protein heterogeneity in the context of the TME, and controls response to BH3-mimetics is unknown and has the potential unlock the deployment of targeted therapies such as BH3-mimetics and NF-κΒ inhibitors (Figure 1).

**Figure 1.**
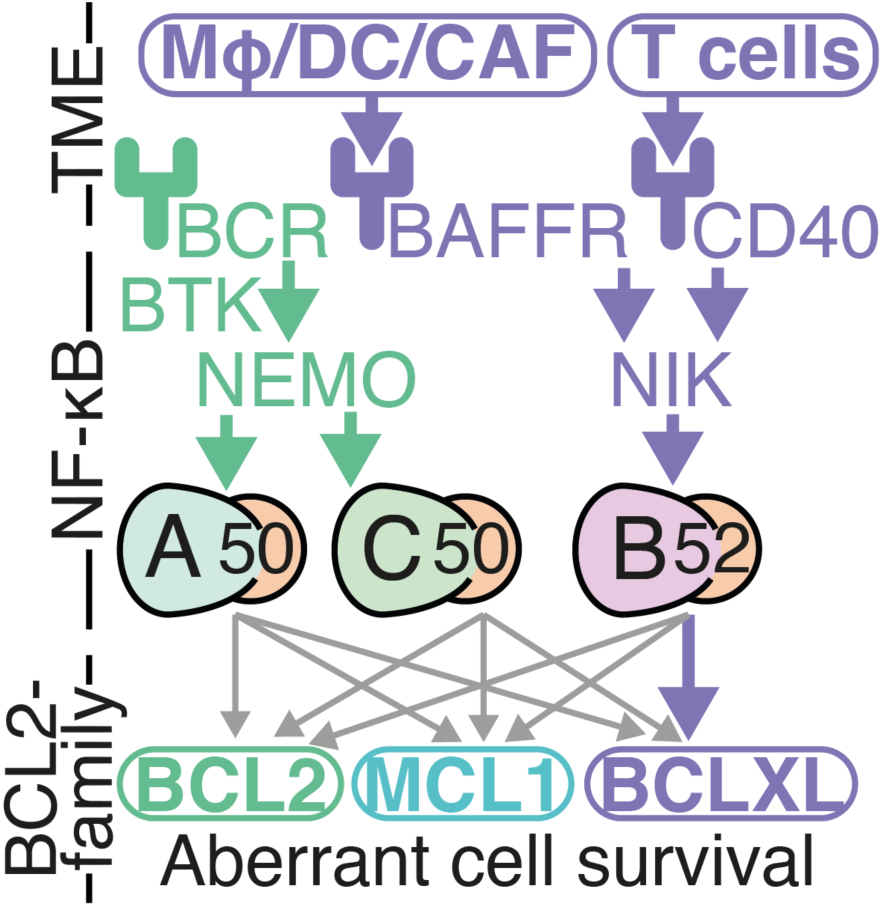
Summary of existing knowledge of links between the Tumour Microenvironment (TME), NF-κB dimers, and BCL2-family proteins in lymphoma. The canonical NF-κB pathway (green) is mediated in part by B-cell receptor (BCR) signaling. BCR signaling transduces signals through Bruton’s tyrosine kinase (BTK), and NF-κB essential modulator (NEMO), to primarily activate RelA- and cRel-containing NF-κB dimers. Single cell RNA sequencing has revealed that the primary impact of the TME on DLBCL cells is through the non-canonical NF-κB pathway (purple). Macrophages (Mϕ), dendritic cells (DC), and cancer associated fibroblasts (CAFS), activate non-canonical signaling through B cell-activating factor receptor (BAFFR). T cells in the TME secrete CD40 ligand. Both BAFFR and CD40 activation lead to stabilization of NF-κB-inducing kinase (NIK) and activation of RelB-containing NF-κB dimers. While RelB activity has been shown to upregulate BCLXL in chronic lymphocytic leukemia, and there are frequently reported links between NF-κB and BCL2, it is unknown which NF-κB dimers induce which BCL2-family proteins (indicated by gray arrows), and how this drives response to therapies in the context of the DLBCL TME.

## Methods

### Mice

Primary B cells were acquired from C57BL/6 WT/IκBε^−/−^/IκBα^mut^ female mice (9-12 weeks old), spleens were homogenized and splenocytes were stained with anti-CD43, prior to purification by negative column selection (Miltenyi Biotec, Cologne Germany) as described (14, 42). B cell purity was assessed by flow cytometry staining for anti-B220. Mice were maintained in environmental control facilities at the University of California, Los Angeles. Animal work was performed according to University of California, Los Angeles regulations, under an approved protocol.

### Cell culture

DLBCL cell lines were sourced from DSMZ (Braunschweig, Germany) and cultured in recommended medium. Cells were kept at 37°C in 5% CO_2_ conditions and routinely tested for mycoplasma contamination. NIH3T3 cells transfected to stably express human CD40 (hCD40L-3T3) were obtained from Prof Chris Pepper (Brighton and Sussex Medical School, UK) and transfection was maintained by puromycin selection (0.1%).

### Co-culture

DLBCL cells were co-cultured with hCD40L-3T3, or non-transfected control cells for 24 hours on 96-well flat bottom plates, with or without small molecule inhibitors.

### Drug Treatments

0.1 x 10^5^ DLBCL cells were plated per well and treated with Amgen16 used at concentrations of 1.25-100μΜ purchased from Merck (Gillingham, UK). ABT199 (venetoclax) was purchased from Selleckchem (Texas, USA) and used at concentrations of 0.0001-100μM with or without Amgen16 at a dose of 50μM for 24 hours.

### Flow Cytometry

Early apoptosis was determined by flow cytometry following Annexin V staining (BioLegend) and data was acquired on a CytoFlex LX, using the BD fix and perm kit (BioLegend), and data was acquired as previously described (19).

### Proteomics

Equal amounts of protein determined using BCA assay, from each sample were digested using trypsin. For tyrosine phosphoproteomics, peptides were enriched using SH2-superbinder beads as described (43), and the flow-through reconstituted in freshly prepared sample loading buffer and subjected to phosphopeptide enrichment using TiO₂ beads preconditioned with the loading buffer for 5 minutes. Data-independent acquisition mass spectrometry (DIA-MS) experiments were performed on a ThermoFisher Orbitrap Astral mass spectrometer. A DIA spectral library generated using DIA-MS2pep was utilized in DIA-NN software for protein quantification, with precursor and protein false discovery rates (FDR) set to 1%. Heatmaps were generated using R package pheatmap (version 1.0.12).

### Immunofluorescent Microscopy

NF-κB activity was determined post fixation with 4% PFA and permeabilization with 0.1% Triton-X. Cells were stained with anti-p65 (Invitrogen), RelB (Cell signaling), cRel (Invitrogen), DAPI (ThermoFisher) and rhodamine phalloidin (Thermo Scientific). Images were captured with *Operetta CLS*, and nuclear:cytoplasmic ratios of NF-κB proteins were analyzed with *Cell Profiler*.

### Simulations

The computational model and simulation method is described previously (40). Basal canonical pathway activation was increasing 0% to 0.5%. NIK degradation was decreased using the equation *NIK deg=1/(1+(t/600))*. 25 individual simulations were run for each cell line with parameter sampling as described previously (40, 44). Models were solved using the Julia DifferentialEquations.jl package (45).

### Statistical Analysis

All statistical analysis was performed using GraphPad Prism 10.0 (California, USA) and Julia 1.8.5. Analysis of paired datasets was performed with paired t-tests, whereas unpaired t-tests and one-way ANOVA to analyze differences between groups, followed by indicated post-hoc tests. LC_50_ values were generated a post four-parameter linear regression test. Flow cytometry data was analyzed on CytExpert Software (Beckman Coulter). \**P*<0.5; \*\**P*<0.01; \*\*\**P*<0.001; \*\*\*\**P*<0.0001.

## Results

### Flow cytometry-based ‘fingerprinting’ predicts sensitivity to inhibitors targeting non-canonical NF-κΒ and BCL2

Previous work in which NF-κB subunit abundance was visualized as contoured density plots standardized using z-scores (Figure 2A) established heterogeneity in RelA abundance between ABC-DLCBL cell lines (19). To establish whether similar variability is present in ABC- and GC-DLBCL we measured the levels of NF-κB in two ABC-(RIVA and U2932) and two GC-DLBCL (SUDHL8 and SUDHL10) cell lines (Figure 2B), and identified varied expression of both RelA and RelB, which did not align with COO (Figure 2B). While there was no variability in RelB between ABC cell lines (RIVA and U2932, Figure 2C), as reported previously (19), GC-DLBCL cell line SUDHL10 was found to have significantly higher RelB (Figure 2B and C) and a higher ratio of RelB to RelA (Figure 2B). We hypothesized that this cell line may be sensitive to NIK inhibition, and found NIK-inhibitor Amgen16 was significantly more effective at killing SUDHL10 cells than cell lines with lower levels of RelB (Figure 2D and E, p<0.01).

**Figure 2.**
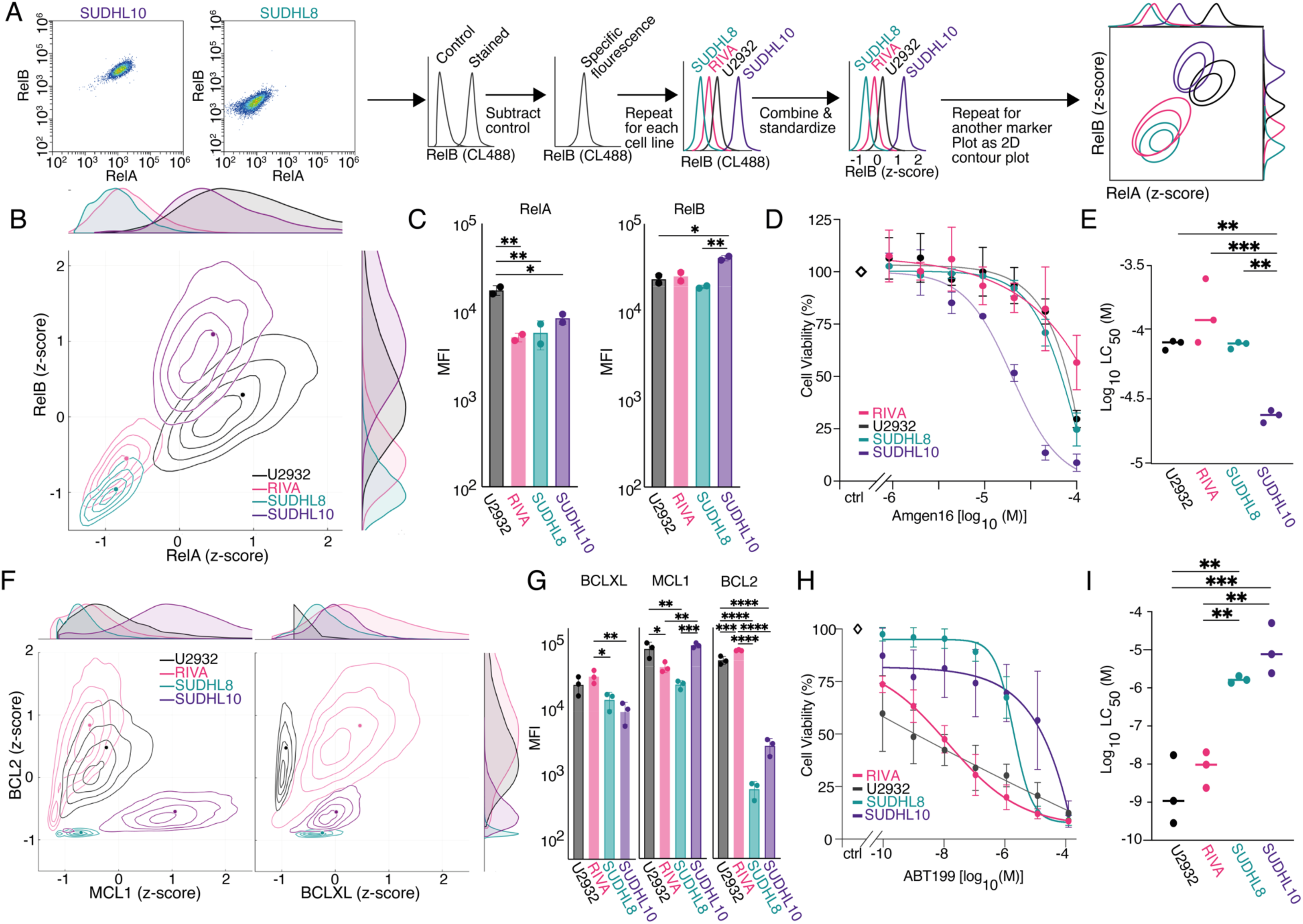
NF-κB and BCL2 ‘fingerprinting’ predicts sensitivities to inhibition of non-canonical NF-κΒ and BCL2 in Diffuse Large B cell Lymphoma (DLBCL). (A) Representative dot plots of RelA and RelB Median Fluorescence Intensity (MFI) values for the DLBCL cell lines SUDHL8, SUDHL10, U2932, and RIVA. Workflow pipeline for the productions of NF-κΒ fingerprints. (B) Relative RelA and RelB expression levels measured with flow cytometry, in the cell lines RIVA, U2932, SUDHL8 and SUDHL10 as fingerprints, with dots representing the median in a single experiment, also shown as probability density curves (right and top). (C) Bar graphs of RelA and RelB median fluorescence intensity (MFI) values as mean ± standard deviation of two independent experiments. (\**P*< 0.05; one-way ANOVA with Tukey’s comparisons test). (D) Cell viability in response to 0.125-100μΜ of Amgen16 in the indicated DLBCL cell lines post a 24-hour treatment, error bars representing the mean ± standard deviation of three independent experiments, normalized to the untreated control. (E) LC_50_ values for the indicated cell lines in response to Amgen16 with mean indicated, three independent experiments *(**P<0.01, ***P<0.001 unpaired T-test).* (F) Relative MCL1, and BCL2, and BCLXL expression levels measured with flow cytometry, in the cell lines RIVA, U2932, SUDHL8 and SUDHL10 as fingerprints, with dots representing the median in a single experiment, also shown as probability density curves (right and top). (G) Bar graphs of MCL1, and BCL2, and BCLXL median fluorescence intensity (MFI) values as mean ± standard deviation of three independent experiments. *(*P<0.05, **P< 0.01, ***P<0.001, ****P<0.0001; one-way ANOVA with Tukey’s comparisons test*). (H) Cell viability in response to 0.0001-100μΜ of ABT199 (venetoclax) in the indicated DLBCL cell lines, post a 24-hour treatment, error bars representing the mean ± standard deviation of three independent experiments, normalized to the untreated control. (I) LC_50_ values for the indicated cell lines in response to ABT199 with mean indicated, of three independent experiments *(**P<0.01, ***P<0.001 unpaired T-test)*.

We next sought to establish whether a similar approach could enable predictable targeting of BCL2-family proteins. We identified two lines with low BCL2 (SUDHL8, and SUDHL10, Figure 2F), and two lines with significantly higher BCL2 (U2932, and RIVA, Figure 2G, p<0.0001). In agreement with clinical observations (31), we found that ABT199 had significantly higher efficacy in cell lines with high BCL2 (21H and I, p<0.01). Previous BH3-profiling had suggested a complete block of apoptosis in SUDHL10 cells due to loss of effector proteins (Bak/Bax) (36); however, multiple recent studies have found substantial Bax expression in SUDHL10 cells, and implicated low BCL2 expression in resistance to venetoclax in SUDHL10 cells (34, 46). NF-κB profiling (Figure 2B) revealed a vulnerability to NIK inhibition (Figure 2D-E), indicating the potential for combined NF-κB and BCL2 “fingerprinting’’ to inform the rational assignment of therapeutic agents targeting multiple pathways in DLBCL.

### BCL2 family protein heterogeneity confers heterogenous responses to ABT1GG within a biclonal DLBCL cell line

The U2932 cell line harbors two phenotypically and genetically distinct subclones. The subclone R1 is CD20, CD38 high, while the R2 subclone is CD20, CD38 low (19, 47). We found that the R1

subclone (Figure 3A) was ABT199 resistant, whereas the R2 subclone was ABT199 sensitive, even at the lowest dose of ABT199 tested (0.0001μΜ) (Figure 3B). While BCL2 protein abundances could not explain the different sensitivities to ABT199 (Figure 3C), we identified significantly higher MCL1 in the R1 subclone (Figure 3C, p<0.0001). To determine if the selective sensitivity of the R2 subclone to ABT199 would be retained when DLBCL cells are activated by their TME we employed a co-culture system of hCD40L-3T3, which multiple studies have shown mimics both cell-contact and the presence of activated T cells in the DLBCL TME (24, 33, 34, 48)(Figure 3D). The R2 subclone became substantially more resistant in the co-culture system (Figure 3E), significantly increasing resistance to ABT199 (Figure 3F, p<0.01). In hCD40L-3T3 BCLXL was significantly more induced than both BCL2 (p<0.05) and MCL1 (p<0.01) which were either unchanged (BCL2) or moderately decreased (MCL1) (Figure 3G). Within the U2932 cell lines we identified multiple distinct resistance mechanisms to ABT199; high MCL1 in the R1 subclone, and high BCLXL in the context of CD40L co-culture.

**Figure 3.**
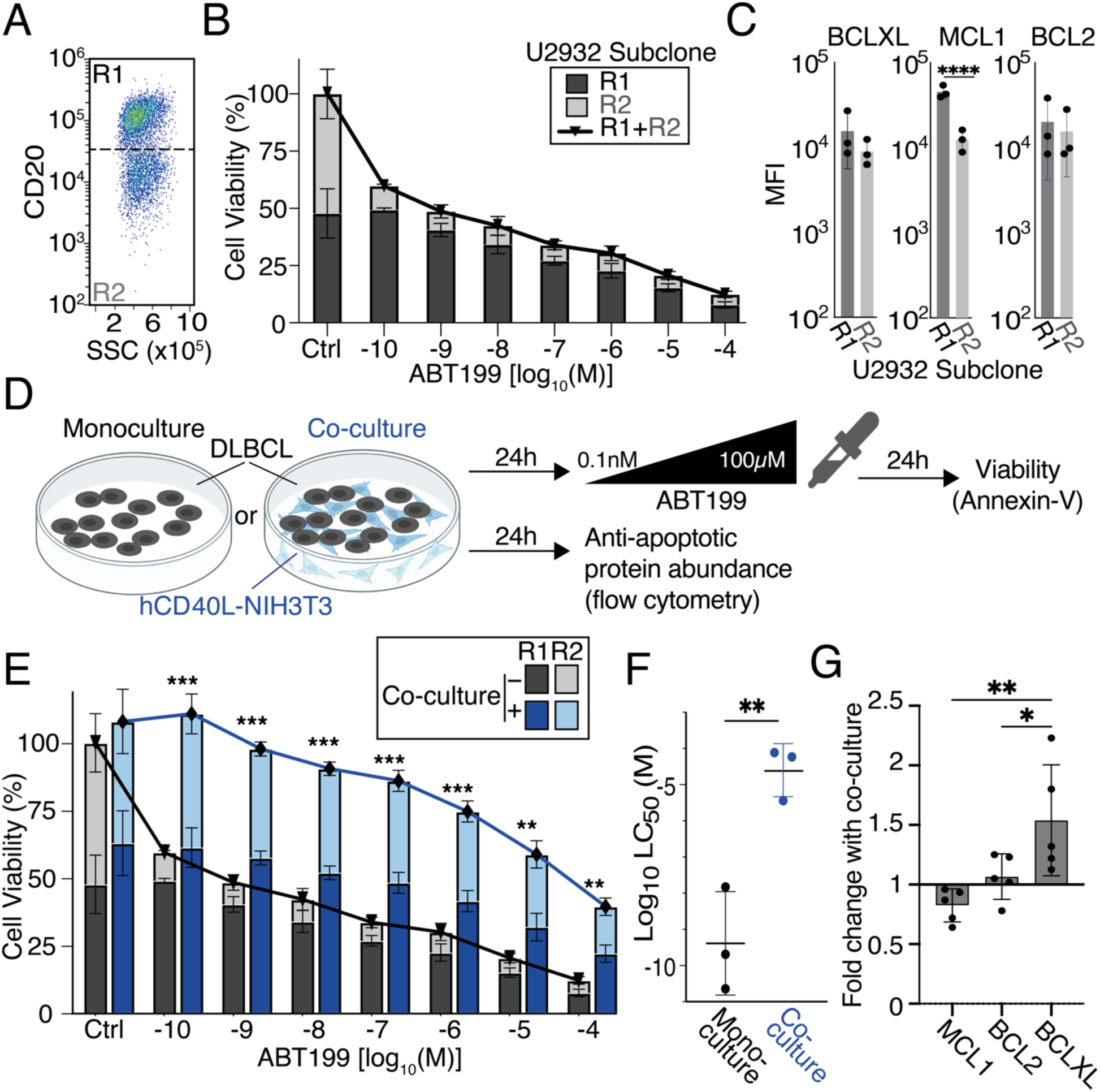
Differential BCL2 family abundance in the sub-clonal cell line U2932 predicts differential responses to ABT199 (venetoclax), TME mimicking hCD40L-3T3 co-culture confers resistance to BCL2 inhibition. (A) Flow cytometry dot plot showing the distinct levels of CD20 Median Fluorescence Intensity (MFI) in the U2932 subclones R1 and R2. (B) Cell viability in response to 0.0001-100μΜ of ABT199in the indicated U2932 subclones R1 and R2, post 24-hour treatment, error bars representing the mean ± standard deviation of three independent experiments, normalized to the untreated control. (C) Bar graphs of MCL1, and BCL2, and BCLXL median fluorescence intensity (MFI) values as mean ± standard deviation of three independent experiments (\*\*\*\**P*<0.0001; unpaired t-test). (C) Cell viability in response to 0.0001-100μΜ of ABT199in the indicated U2932 subclones R1 and R2, post 24-hour treatment, error bars representing the mean ± standard deviation of three independent experiments, normalized to the untreated control. (D) Schematic of the co-culture system used. U2932 cells were co-cultured with hCD40L-3T3 cells prior to treatment with increasing doses of ABT199. (E) Viability of U2932 cells cultured for 24 hours with or without CD40-Ligand NIH3T3 in the presence of 0.0001-100μΜ of ABT199. Error bars representing the mean ± standard deviation of three independent experiments, normalized to the untreated control *(**P<0.01, ***P<0.001 multiple unpaired T-tests using the Holm-Šídák corrections test)*. (F) LC50 values for the U2932 cell line in monoculture vs co-culture with hCD40L-3T3 in response to ABT199, with error bars representing the mean ± standard deviation of three independent experiments*(**P<0.01, unpaired T-test). (G)* MCL1, BCL2, and BCLXL levels, shown as fold changes of 24-hour stimulation with hCD40L-3T3 cells to unstimulated control in U2932 cells with error bars representing the mean ± standard deviation of five independent experiments *(*P<0.05, **P<0.01 one-way ANOVA with Tukey’s comparisons test)*.

### TME-mediated resistance to BH3-mimetics can result from induction of multiple compensatory BCL2-family proteins

Repeating analysis in the BCL2-dependent RIVA cells identified significant TME-mediated ABT199 resistance in hCD40L-3T3 co-culture (Figure 4A and B, p<0.05) and a significant increase in BCLXL (Figure 4C, p<0.01). This effect was dependent on CD40L as untransfected NIH3T3s did not upregulate BCLXL (supplemental Figure 1,2 and 4).

**Figure 4.**
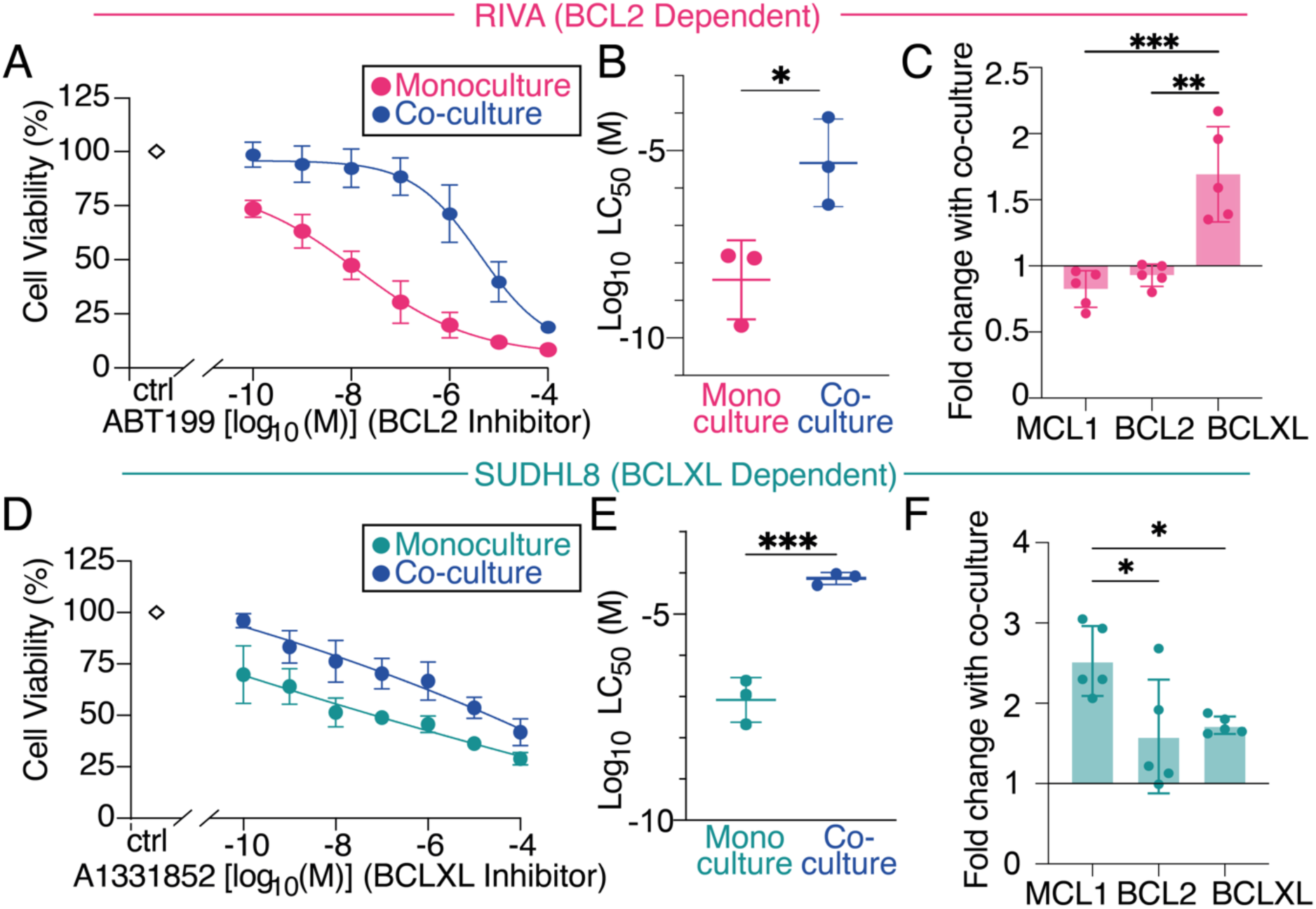
Tumor microenvironment-mimicking co-culture induced resistance to BH3-mimetics in Diffuse Large B Cell Lymphoma (DLBCL) cell lines. (A) Viability of RIVA cells cultured for 24 hours with or without hCD40L-3T3 cells in the presence of 0.0001-100μΜ of the BCL-2 inhibitor ABT199 (venetoclax). Error bars representing the mean ± standard deviation of three independent experiments, normalized to the untreated control. (B) LC_50_ values for RIVA in each condition with error bars representing the mean ± standard deviation of three independent experiments, normalized to the untreated control (*\*P<0.05*, unpaired T-test). (C) MCL1, BCL2, and BCLXL levels, shown as fold changes of 24-hour stimulation with hCD40L-3T3 cells to unstimulated control in RIVA with error bars representing the mean ± standard deviation of five independent experiments *(**P<0.01, ***P<0.001* one-way ANOVA with Tukey’s comparisons test. (D) Viability of SUDHL8 cells cultured for 24 hours with or without hCD40L-3T3 cells in the presence of 0.0001-100μΜ of the BCLXL inhibitor A1331852. Error bars representing the mean ± standard deviation of three independent experiments, normalized to the untreated control. (E) LC_50_ values for SUDHL8 in each condition with error bars representing the mean ± standard deviation of three independent experiments, normalized to the untreated control (*\*\*\*P<0.001*, unpaired T-test). (F) MCL1, BCL2, and BCLXL levels, shown as fold changes of 24-hour stimulation with hCD40L-3T3 cells to unstimulated control in SUDHL8 with error bars representing the mean ± standard deviation of five independent experiments *(*P<0.05,* one-way ANOVA with Tukey’s comparisons test).

To establish whether this effect was unique to venetoclax we employed the SUDHL8 cell line, which has established sensitivity to BCLXL inhibition by the BH3-mimetic A1331852 (34). SUDHL8 cells were significantly resistant to BCLXL inhibition in hCD40L-3T3 co-culture (Figure 4D-E, p<0.001). Significantly more MCL1 was induced than either BCLXL or BCL2 in hCD40L-3T3 co-culture (Figure 4F, p<0.05). This effect was not seen in co-culture with untransfected NIH3T3s (supplemental Figure 3 and 4). While CD40 is known to activate non-canonical NF-κB signaling component RelB, which has been shown to control BCLXL expression in related hematological malignancies (25), the control of MCL1 by specific NF-κB subunits is less clear, particular in the context of lymphoma (Figure 1A).

### The SUDHL8 cell line has high basal nuclear NF-κΒ RelA activity through elevated B cell receptor signaling

We hypothesized that the canonical RelA- or cRel-containing NF-κB dimers could be responsible for cell line-specific induction of BH3-mimetic resistance through MCL1 (Figure 5A). Quantification of the nuclear activity of NF-κB RelA and cRel using immunofluorescence microscopy (Figure 5B, Supplemental Figure 5) revealed substantial cell-to-cell variability in nuclear NF-κB activity (Figure 5C), and significantly higher nuclear RelA in the SUDHL8 cell line (Figure 5D, bottom).

**Figure 5.**
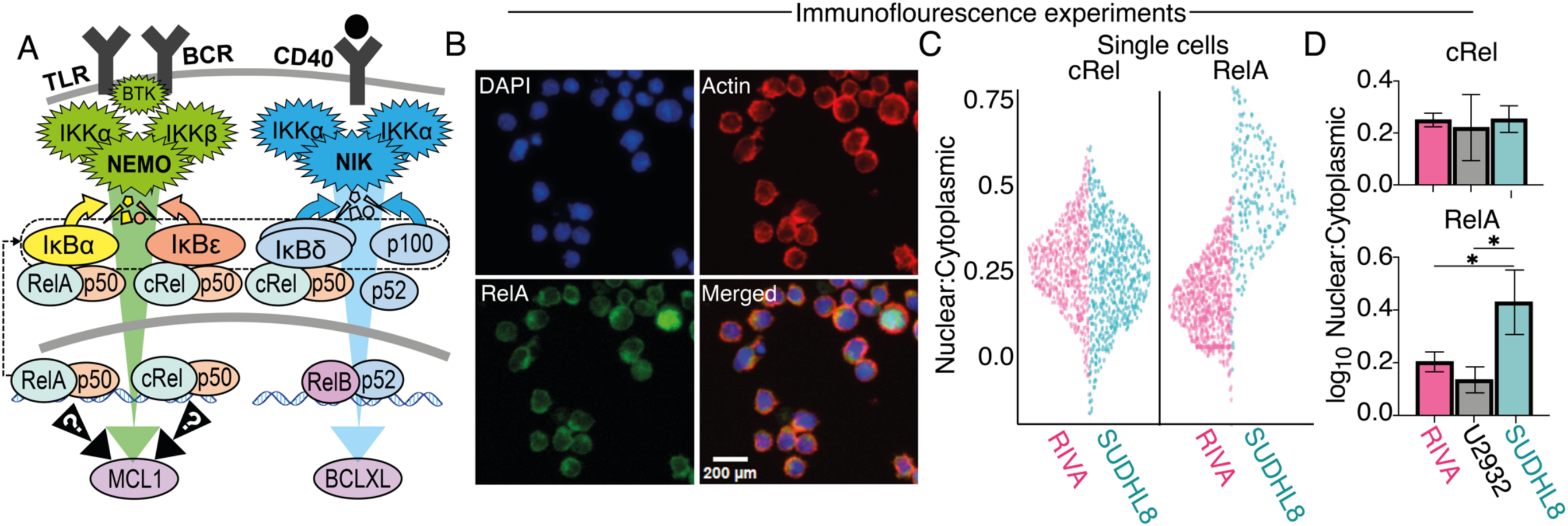
Immunofluorescence microscopy reveals differential NF-κB basal RelA activity in Diffuse Large B-cell Lymphoma (DLBCL) cell lines. (A) Schematic of the canonical and non-canonical NF-κB signaling pathway illustrating how non-canonical NF-κB activation results in the induction of BCLXL and how elevated basal canonical signaling induces multiple negative regulators of NF-κB which may contribute to cell line specific induction of MCL1. (B) Representative immunofluorescence microscopy following staining in RIVA cells for RelA (NF-κB), DAPI (nucleus), actin (cytoplasm), to measure nuclear and cytoplasmic subcellular localization. (C) Quantification immunofluorescence microscopy showing nuclear to cytoplasmic ratio of cRel and RelA in single cells. Data shows a representative replicate in the indicated cell lines. Violin width indicates data density. (D) Quantification of the nuclear to cytoplasmic ratio of cRel (top) and RelA (bottom) as bar graphs with error bars of the mean ± standard deviation of three independent experiments in RIVA, U2932 and SUDHL8 (*P<0.05; One-Way ANOVA).

We performed global, and serine threonine proteomics in the cell line with elevated nuclear RelA (SUDHL8), a line with low nuclear RelA (U2932), and a line dependent on non-canonical signaling (SUDHL10), to test whether elevated B Cell Receptor (BCR) signaling was responsible for elevated nuclear RelA. These data show substantially elevated CD19 and SYK abundance and phosphorylation levels in SUDHL8 compared to U2932 and SUDHL10 (supplemental Figure 6A-B). BTK abundance in SUDHL8 was not elevated; however, phosphorylated Y223 in the BTK SH3 domain has recently been found to mirror catalytic activity (49). Phosphotyrosine (pY) proteomics revealed elevated phosphorylation at BTK_Y223 in SUDHL8 cells (1.8 fold) consistent with elevated BTK activity (supplemental Figure 6C).

As CD40L is a selective activator of non-canonical NF-κΒ signaling in B cells (50), it is not clear how elevated BTK activity and RelA nuclear abundance in SUDHL8 could confer CD40-mediated induction of MCL1 (Figure 5A). IκBα, IκBε, and p100/IκBδ are all induced by RelA activity (12), and are preferential inhibitors of distinct NF-κB subunits (Figure 5A)(14, 40, 42). Therefore, we hypothesized that the altered profile of NF-κΒ inhibitors could control distinct responses to the TME, and sought to establish how this could lead to selective induction of MCL1.

### Computational modelling informed by imaging experiments predicts that high basal nuclear NF-κΒ RelA leads to crosstalk, with CD40-mediatead induction of cRel

Employing established computational models of NF-κB signaling (19, 40), and running two simulations differing only in the degree to which basal BCR-signaling was elevated, predicted that RelA:p50 is mainly inhibited by IκBα in RIVA cells (Figure 6A). SUDHL8 cells were predicted to have reduced inhibition of RelA and a moderate shift in inhibitor composition (away from IκBα and IκBε, Figure 6A–left). The predicted impact of high basal canonical pathway activation on cRel:p50 was markedly different (Figure 6A, right), shifting towards IκBδ inhibition of cRel in SUDHL8 cells, with <10% of cRel being inhibited by IκΒα and -ε (Figure 6A right).

**Figure 6.**
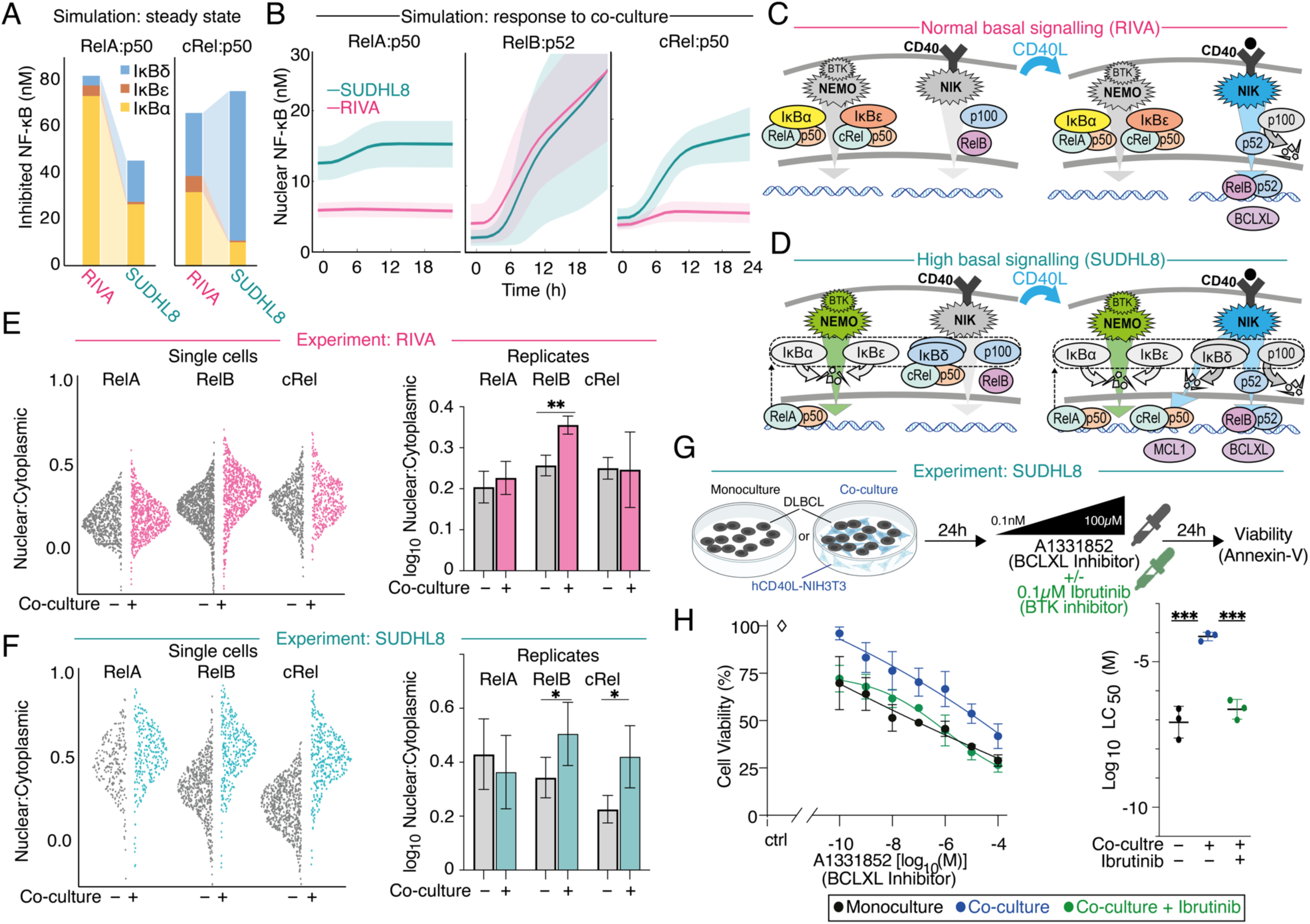
Co-culture with hCD40L-3T3 cells upregulates NF-κB RelB and BCLXL in DLBCL, and selectively upregulates NF-κB cRel and MCL1 through signaling crosstalk when NF-κB RelA is chronically active. (A) Abundances of the heterodimers RelA:p50 (left) and cRel:p50 (right) bound to inhibiting NF-κΒ proteins (IκB), as simulated by computational modeling for basal canonical NF-κΒ activation state in RIVA, and high basal activation state in SUDHL8. (B) Computational modeling results showing the nuclear abundance of the indicated NF-κB dimers in a simulation with low basal nuclear RelA (RIVA, pink), and a simulation high basal nuclear RelA (SUDHL8, teal). The mean (line) and standard deviation (shaded region) of 25 cells is indicated. (C) Schematic demonstrating the mechanism by which NF-κB contributes to the induction of BCLXL in a cell line with normal basal signaling (RIVA) upon stimulation with CD40L (right). hCD40L-3T3 mediated activation of NIK results in p100 processing into p52, leading to nuclear translocation of RelB:p52 and increased expression of BCLXL. Gray = inactive pathways and low abundance proteins, color = active pathways and predominant complexes. (D) Schematic demonstrating how crosstalk emerges between CD40, NIK, cRel, and MCL1. Increase basal RelA results in increase p100, and IκBδ (left). hCD40L-3T3 mediated activation of NIK results in processing of IκBδ and release of cRel:p50, which translocates to the nucleus and potentially upregulates expression of MCL1 (right). Gray = inactive pathways and low abundance proteins, color = active pathways and predominant complexes. (E-F) Quantification of immunofluorescence microscopy in RIVA (E) and SUDHL8 (F) cells showing nuclear to cytoplasmic ratio of RelA, RelB and cRel. Data post 24 hours of monoculture and hCD40L-3T3co-culture is shown side by side. Single cells of a representative replicate are shown (left), with violin width indicating data density. Quantification of the nuclear to cytoplasmic ratio of RelA, RelB and cRel is shown as bar graphs (right) with error bars displaying the mean ± standard deviation of three independent experiments (*\*P<0.05, **P<0.01*; unpaired t-test). (G) Schematic of the co-culture system used, following 24 hours of treatment with 0.0001-100μΜ of the BCLXL inhibitor A1331852 ± 0.1μM of the Bruton’s Kinase (BTK) inhibitor Ibrutinib. (H) Cell viability of SUDHL8 in response to 0.0001-100μΜ of the BCLXL inhibitor A1331852 post a 24-hour treatment in monoculture, post a 24-hour hCD40L-3T3co-culture or post a 24-hour hCD40L-3T3 co-culture with the addition of 0.1μM of the BTK inhibitor, Ibrutinib. LC_50_ values for each condition is shown (right) with error bars representing the mean ± standard deviation of three independent experiments, normalized to the untreated control (*\*\*\*P<0.001* one-way ANOVA with Tukey’s comparisons test).

The steady state of these simulations (Figure 6B, t=0) recapitulated the selectively elevated nuclear RelA, seen in immunofluorescence imaging (Figure 5D). Simulating co-culture mediated non-canonical pathway activation revealed induction of RelB:p52, without induction of RelA:p50 (Figure 6B, left and center). However, induction of nuclear cRel:p50 was predicted only in the SUDHL8 cell line (Figure 6B, right). Cells with low basal nuclear RelA, such as RIVA cells, reliably induce p100 processing to p52, and subsequent translocation of RelB:p52 to the nucleus upon CD40L activation (Figure 6C). Simulations predict that constitutive activation of the BCR pathway results in degradation of IκBα and IκBε, nuclear translocation of RelA:p50 and subsequent induction of p100, which forms higher inhibitory complexes termed IκBδ (13). IκBδ has higher affinity for cRel than RelA (14), resulting in sequestration of cRel by IκBδ prior to CD40L activation (Figure 6D, left). Upon CD40L activation p100 is processed into a self-inhibited p100:p52 dimer, which frees cRel for nuclear translocation (13, 42) (Figure 6D, right). This crosstalk mechanism was first identified in fibroblasts (13), and shown to preferentially upregulate cRel in proliferating healthy B cells responding to BAFF (42). This predicted crosstalk from non-canonical signaling to NF-κB cRel has not previously been described in lymphoma or linked to induction of anti-apoptotic proteins, therefore we sought to validate these computational predictions.

### Experimental validation of modelling predictions confirms crosstalk between CD40 and cRel confers BH3-mimetic resistance and can be overcome with BTK inhibition

Immunofluorescence imaging confirmed that hCD40L co-culture did not induce RelA (Figure 6E-F, compare to 6B left), but did induce significantly higher levels of nuclear:cytoplasmic RelB in both the RIVA and SUDHL8 cell lines, as predicted by computational simulations (Figure 6E-F, compared to 6B middle, p<0.05). Only in SUDHL8 cells was this accompanied by significantly increased nuclear cRel, as predicted computationally (Figure 6E-F, compared to 6B right, p<0.01). hCD40L co-culture with the addition of ibrutinib (targeting BTK) re-sensitized SUDHL8 cells to BCLXL inhibition, confirming BCR-pathway activation was responsible for BH3-mimetic resistance in the SUDHL8 cells (Figure 6G-H, p<0.001).

### NF-κΒ cRel activity selectively induces MCL1 in B cells

We hypothesized that crosstalk-mediated induction of cRel was responsible for the induction of MCL1 (Figure 4F). ChIP-seq data generated in a lymphoblastoid B cell line (GM12878)(51), showed promiscuous binding of all NF-κB subunits around the transcription start site (TSS) of *BCL2L1* (encoding BCLXL), and no definitive binding of NF-κB subunits at the TSS for *BCL2*. The same data revealed marginally higher levels of cRel binding than both RelB and RelA at the *MCL1* TSS (Figure 7A). To explore whether this was unique to the lymphoblastoid cell lines we collected Model-based Analysis for ChIP-Seq 2 (MACS2) scores for NF-κB cRel, RelA and RelB within 1kb of the MCL1 TSS across ChIP-Seq datasets reported in a ChIP-seq database (ChIP-Atlas (52), Figure 7B) confirmed significantly higher binding of cRel than both RelA and RelB (Figure 7B, p<0.0001).

**Figure 7.**
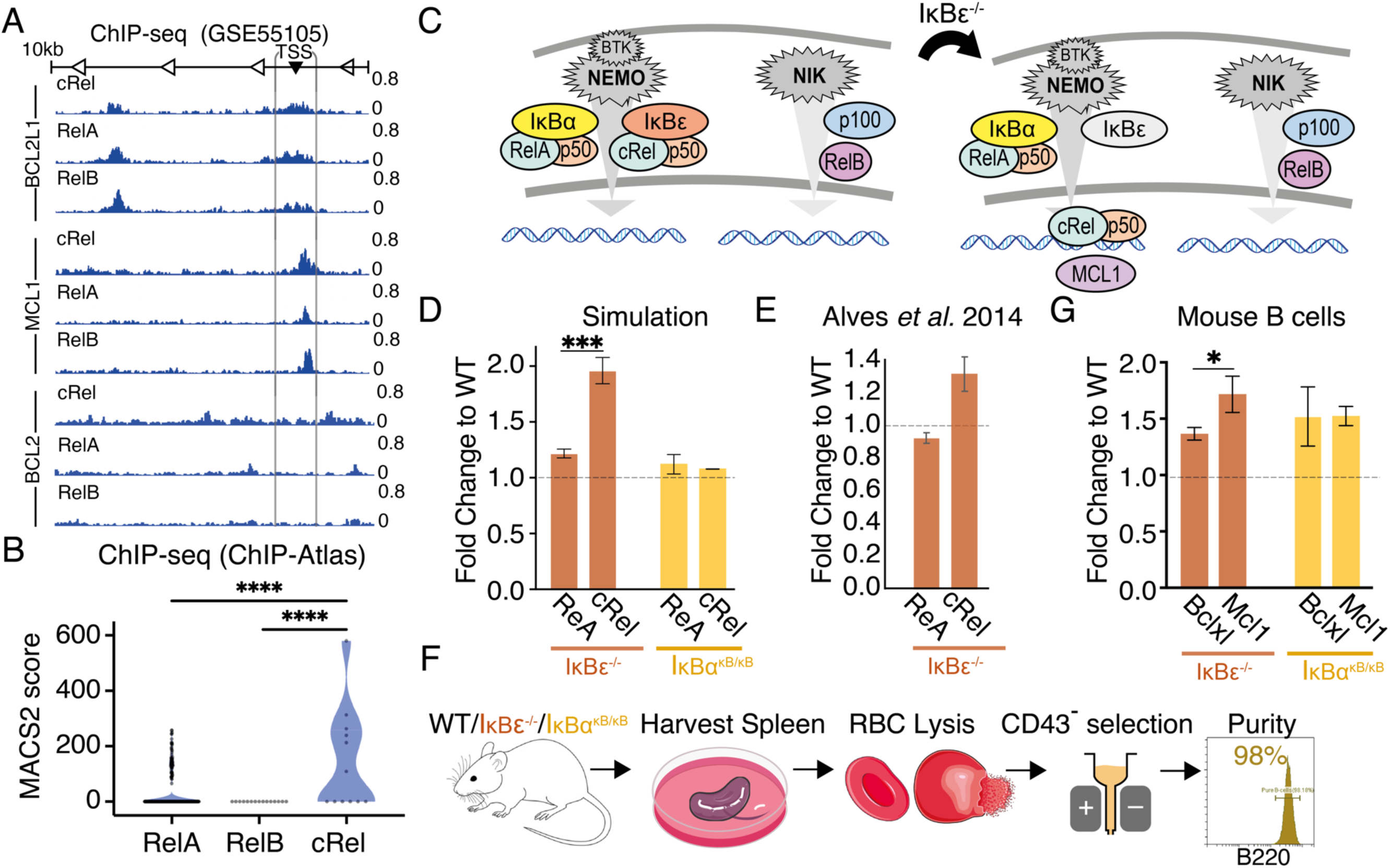
IκBε knock-out mouse model reveals cRel-dependent induction of MCL1. (A) NF-κB ChIP-seq data from Zhao et al. 2014 (52) showing binding of the indicated NF-κB subunits around the transcription start site (TSS) of the indicated anti-apoptotic genes in lymphoblastoid B cell line. A 1kb window around the TSS is indicated. (B) Model-based Analysis for ChIP-Seq 2 (MACS2) scores for the indicated NF-κB subunits within 1kb of the MCL1 promoter across all ChIP-Seq datasets reported in ChIP Atlas Zou et al. 2024 (52), after outlier removal using ROUT method (q=1%, *\*\*\*\*P<0.0001*). (C) Schematic of normal basal NF-κΒ signaling (left), schematic of NF-κB signaling predicting the unknown effect of IκΒε^-/-^ on cRel and MCL-1 induction. (D) Computational modelling simulation of RelA and cRel abundance in IκBε^-/-^ and IκBα^κB/κB^ demonstrated as fold changes normalized to wild-type. Error bars represent the mean and ± standard deviation of 25 cells *(***P<0.001*, unpaired t-test). (E) RelA and cRel abundances in IκBε^-/-^ and IκBα^κB/κB^ cells derived from primary splenocytes, demonstrated as fold changes to wild-type, error bars representing the mean ± standard deviation. (F) Experimental pipeline showing the workflow of isolating and purifying primary B cells from wild type, IκBε^-^/^-^ and IκBα^κB/κB^ mouse genotypes. (G) Abundances of MCL1 and BCLXL acquired with flow cytometry in primary B cells isolated and purified from IκBε^-^/^-^ (left) and IκBα^κB/κB^ (right) mouse genotypes, demonstrated as fold changes normalized to wild-type. Error bars represent the mean ± standard deviation of three independent experiments (*\*P<0.05*, unpaired t-test).

We sought an alternative method to selectively increase cRel activity in B cells. Computational modelling of NF-κB in wild type (WT) B cells, IκBε knock-out B cells, and B cells bearing an IκBα promoter mutation that lowered IκΒα expression (IκBα^κB/κB^), predicted that IκΒε^-/-^ induces cRel activity significantly more than RelA activity (Figure 7D, p<0.001), while IκBα^κB/κB^ would be non-selective (Figure 7D). This was experimentally validated using data from IκΒε^-/-^ mouse B cells (Alves *et al.* 2014, Figure 7D-E). Primary splenic B cells from IκΒε^-/-^ mice (Figure 7F, 95-99% purity, supplemental Figure 7) had significant selective induction of Mcl1 (Figure 7G, p<0.05), while IκBα^κB/κB^ mice did not (Figure 7G). Together, this ChIP-seq analysis, computational modeling, and validation in primary mouse models (Figure 7D-G) implicate cRel in the upregulation of MCL1 in DLBCL.

### Inhibition of the NF-κB inducing kinase (NIK) overcomes tumor microenvironment (TME)-mediated BH3-mimetic resistance in DLBCL

We sought to identify a single pharmacological intervention that could overcome the multiple methods of TME-mediated resistance to multiple BH3-mimetics identified here. NIK inhibition using Amgen16 (Figure 8A-B) achieved significant re-sensitization of the SUDHL8 cell lines to BCLXL inhibition in hCD40L-3T3 co-culture (Figure 8C, p<0.001). Both cell lines that were ABT199 sensitive in monoculture (U2932 and RIVA), and resistant in hCD40L-3T3 co-culture, were significantly re-sensitized to ABT199 through treatment with the NIK inhibitor (Figure 8E and E, p<0.05).

**Figure 8.**
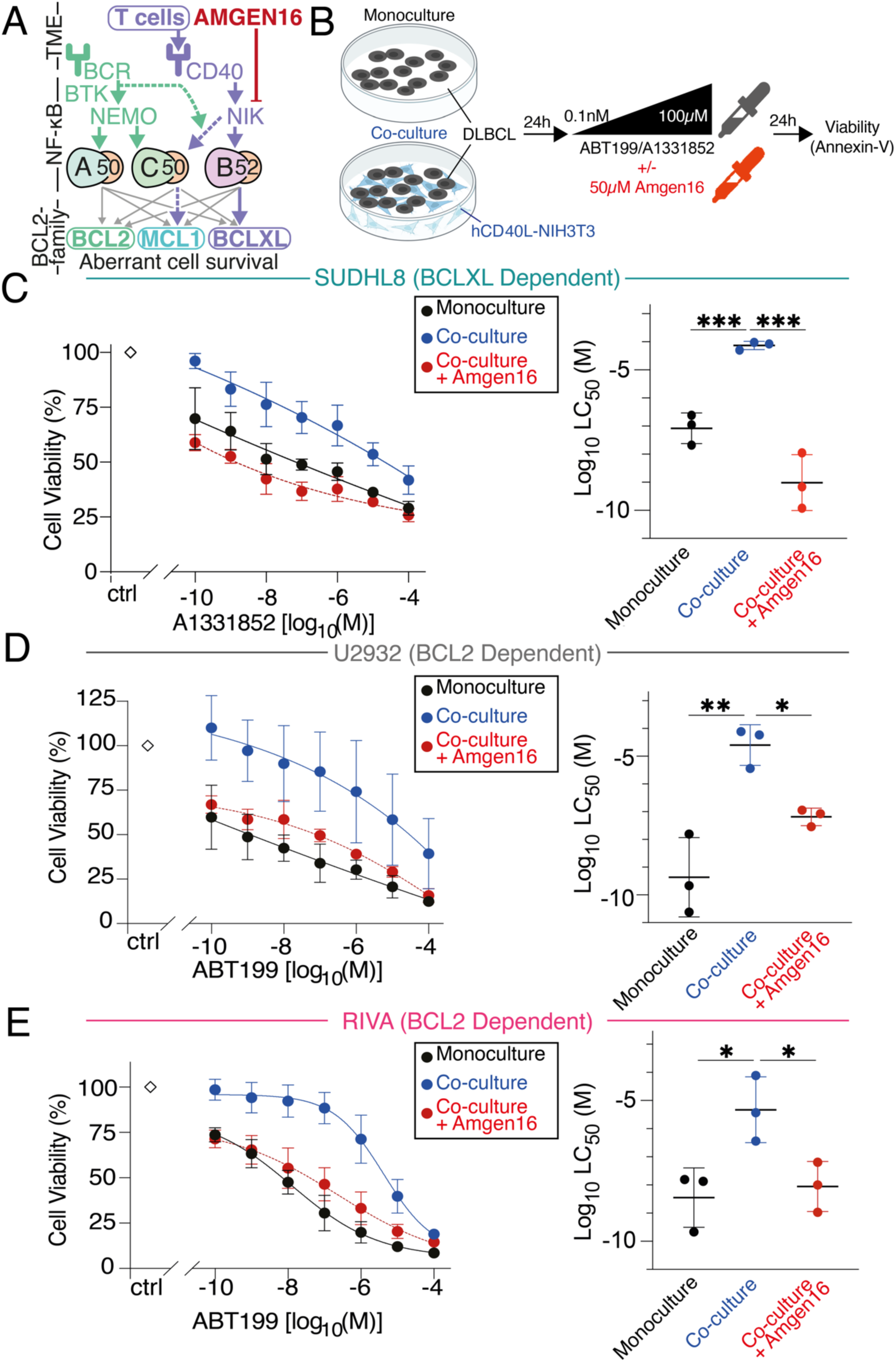
Targeting the NF-κB inducing kinase (NIK) to overcome tumor microenvironment (TME) resistance in Diffuse large B cell lymphoma (DLBCL). (A) Overview of signaling between TME, NF-κB, and BCL2-family proteins indicating interactions revealed here with dashed lines, and NIK-inhibitor Amgen16 displayed in red. (B) Schematic of the co-culture system used, following 24 hours of treatment with 0.0001-100μΜ of venetoclax (ABT199) or A1331852 ± 50μM of the NIK inhibitor Amgen16. (C) Cell viability of SUDHL8 in response to 0.0001-100μΜ of A1331852 post a 24-hour treatment in monoculture, post 24-hour co-culture on hCD40L-3T3 cells or post 24-hour hCD40L-3T3 co-culture with the addition of 50μM of the NIK inhibitor Amgen16. LC_50_ values for each condition is shown (right) with error bars representing the mean ± standard deviation of three independent experiments, normalized to the untreated control (*\*\*\*P<0.001* one-way ANOVA with Tukey’s comparisons test). (D) U2932 and (E) RIVA cells in response to 0.0001-100μΜ of ABT199 in the cell lines post a 24-hour treatment in monoculture, post a 24-hour hCD40L-3T3 co-culture or post 24-hour hCD40L-3T3 co-culture with the addition of 50μM of the NIK inhibitor, Amgen16. LC_50_ values for each condition is shown (right) with error bars representing the mean ± standard deviation of three independent experiments, normalized to the untreated control (*\*P<0.05, **P<0.01* one-way ANOVA with Tukey’s comparisons test.

## Discussion

The molecular heterogeneity that characterizes DLBCL contributes to clinical resistance and relapse (53, 54). While there is substantial evidence that aberrant NF-κΒ activation within the lymph node TME, confers worse prognosis in DLBCL, the pharmacological targeting of this interaction has been challenging due to toxicities associated with targeting the canonical NF-κB pathway (55). The present study indicates that flow cytometry fingerprinting may enable identification and assignment of NIK-inhibitors particularly in the recently identified subset of DLBCL with high RelB and poor prognosis (56). BCL2 fingerprints confirmed that basal high basal BCL2 levels predicted sensitivity to ABT199, in accordance with results from clinical trials (57, 58). When compared to other biochemical or functional measurements such as BH3-profiling flow cytometry-based fingerprinting enables identification of cellular sub populations within a heterogeneous tumor. We found that even within a single cell line we could identify multiple venetoclax resistance mechanisms, including inherent resistance due to high MCL1, and resistance acquired in the context of a TME-mimicking co-culture through upregulation of BCLXL.

BCL2-targeting therapies create impressive responses in CLL, where the bulk of the disease is circulating in the peripheral blood, however lymph-node mediated treatment resistance has been implicated in the inability to cure CLL (59). In DLBCL, where the bulk of the disease is supported by the lymph node TME, BCL2-targeting therapies have been far less effective. Increased non-canonical NF-κΒ activity, and BCLXL, may provide a rationale for the lack of efficacy of ABT199 in DLBCL. NIK inhibition was effective in all cell lines tested here: either as a monotherapy in RelB high DLBCL (Figure 2B), to re-sensitize BCL2-sensitive DLBCL to ABT199 in the TME (Figure 8D and E), or to re-sensitize BCLXL-sensitive DLBCL to BCLXL targeting BH3-mimetics in the TME (Figure 8C). This finding is supported by studies in CLL (24, 25).

We discovered unexpected induction of MCL1 in cells with elevated basal NF-κΒ activity. Incorporating NF-κΒ activity information into computational models led to a prediction of signaling crosstalk between the two NF-κΒ pathways, which was experimentally confirmed. Our data implicate cRel as a regulator of MCL1 in DLBCL, which was previously unknown (60). cRel-mediated gene expression has been correlated with GC-DLBCL (61), and pre-clinical MCL1 inhibitors show increased efficacy in GC-DLBCL (62). As MCL1 inhibitors are currently reaching first-in-human studies (63), a predictive understanding of how the TME controls abundance of anti-apoptotic proteins through NF-κB will be key to successful trial design and clinical adoption. This study highlights the importance of understanding the impact of TME-mediated signaling, which may not be captured in assays performed on isolated cells or isolated mitochondria (64).

Computational simulations can: predict DLBCL responses to BH3-mimetics (39), integrate flow cytometry and mutational information to predict DLBCL signaling responses to the TME (19), and predict unexpected crosstalk between the TME and anti-apoptotic proteins. Integration of high dimensional data with computational modelling enables assignment of targeted therapies that can overcome the protective effect of the TME. While detailed molecular characterization is unlikely to be incorporated into the standard of care for all DLBCL patients, computational simulations informed by genetic profiling of diagnostic samples enable the identification of DLBCL patients that are molecularly predisposed to a poor response (41). Such a prediction motivates further combined molecular profiling and computational modeling in order to assign efficacious targeted therapeutic approaches to the subset patients who will not respond to the standard of care.

## Acknowledgments

We would like to thank all members of the Mitchell and Pepper labs for valuable discussions. This study was supported by Leukaemia UK John Goldman Fellowship 2020/JGF/003, UKRI Future Leaders Fellowship (MR/T04188S/1) and Paul Stanforth PhD Studentship (to S.M), Blood Cancer UK 23004 (to C.P), MRC MR/V00S0S5/1 (to A.P), NIH R01AI132731 and R01AI1278C7 (to A.H).

## Data Availability Statement

The computational code and analysis pipelines generated during and/or analyzed during the current study are available in the GitHub repository, https://github.com/SiFTW/VareliEtAl/. Any other data generated during and/or analyzed during the current study are available from the corresponding author on reasonable request.

## Authorship Contributions

Contribution: **A.V** designed the research, performed experiments, analyzed data, designed figures and wrote the manuscript; **H.V.N** performed experiments, cosupervised part of the study and reviewed the manuscript; **H.C, E.J, L.S** performed experiments; **H.Z., Y.L** performed experiments, analyzed data, designed figures and wrote the manuscript; **F.S** interpreted data and reviewed the manuscript; **A.H** contributed vital new reagents, interpreted data, supervised part of the study and reviewed the manuscript; **A.P, C.P** designed the research, cosupervised the study and reviewed the manuscript; **S.M** designed the research, supervised the study, performed experiments, analyzed data, designed figures and wrote the manuscript.

