## Supplemental Figures for "Systems biology-enabled targeting of NF-κΒ and BCL2 overcomes microenvironment-mediated BH3-mimetic resistance in DLBCL"

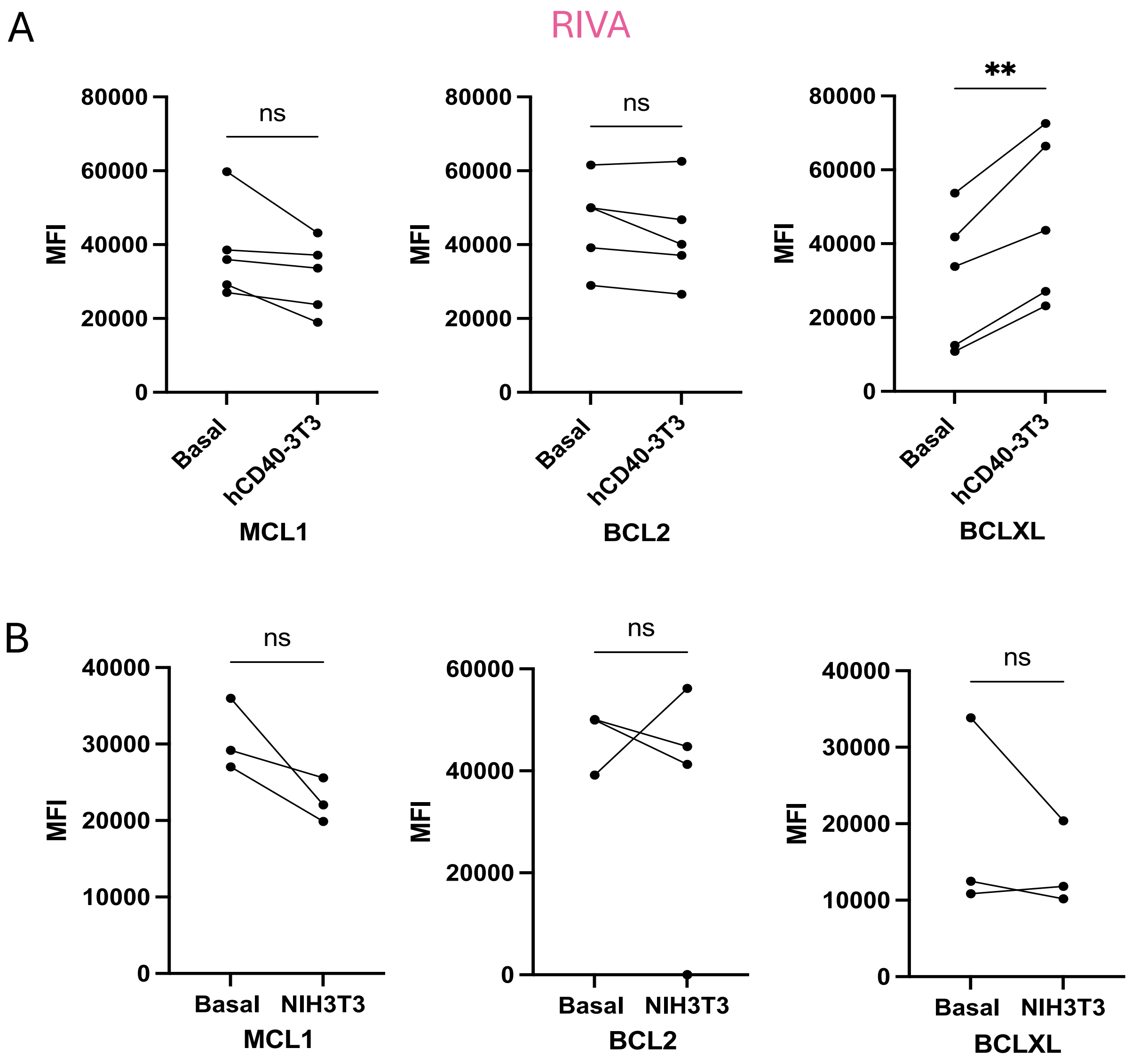


**Supplemental Figure 1.** Higher anti-apoptotic levels are seen in DLBCL cells when are co-cultured with CD40L transfected fibroblasts. (A) RIVA cells were incubated for 24 hours with or without the presence of CD40L transfected fibroblasts. After the stimulation, the DLBCL cells were harvested and stained for Mcl-1, Bcl-2 and Bcl-xL, and then assessed by using flow cytometry. Statistical analysis was performed on five independent experiments' Mean Fluorescence Intensity (MFI) values, between unstimulated and CD40L stimulated RIVA cells, (**$P$<0.01, paired T-test). (B) RIVA cells were incubated for 24 hours with or without the presence of fibroblasts. After the stimulation, the DLBCL cells were harvested and stained for Mcl-1, Bcl-2 and Bcl-xL, and then assessed by using flow cytometry. Statistical analysis was performed on three independent experiments Mean Fluorescence Intensity (MFI) values, between the monocultured and co-cultured RIVA cells, (paired T-test).

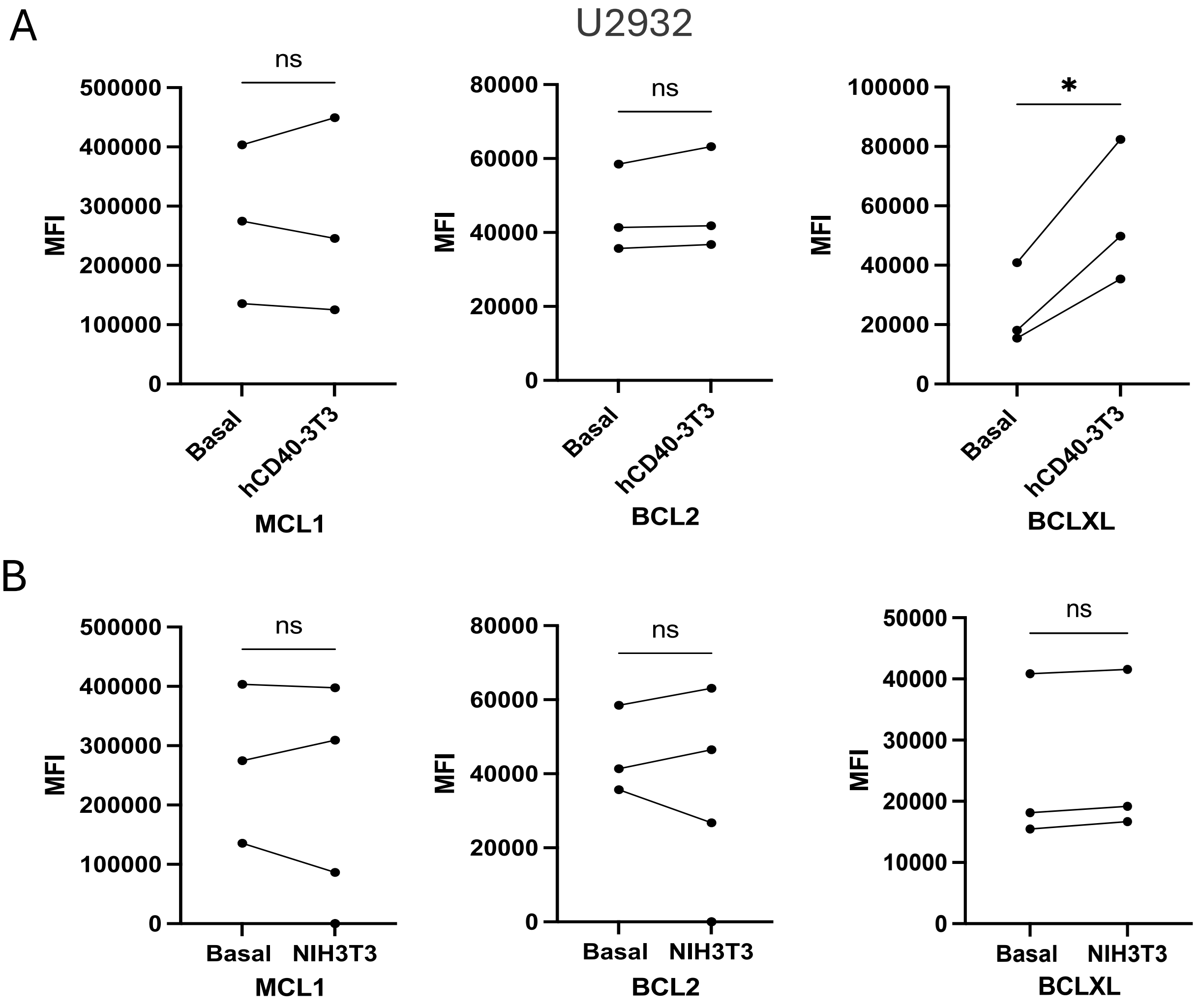


**Supplemental Figure 2.** Higher anti-apoptotic levels are seen in DLBCL cells when are co-cultured with CD40L transfected fibroblasts. (A) U2932 cells were incubated for 24 hours with or without the presence of CD40L transfected fibroblasts. After the stimulation, the DLBCL cells were harvested and stained for Mcl-1, Bcl-2 and Bcl-xL, and then assessed by using flow cytometry. Statistical analysis was performed on three independent experiments' Mean Fluorescence Intensity (MFI) values, between unstimulated and CD40L stimulated U2932 cells, (*$P<0.05$, **$P<0.01$ paired T-test). (B) U2932 cells were incubated for 24 hours with or without the presence of fibroblasts. After the stimulation, the DLBCL cells were harvested and stained for Mcl-1, Bcl-2 and Bcl-xL, and then assessed by using flow cytometry. Statistical analysis was performed on three independent experiments Mean Fluorescence Intensity (MFI) values, between the monocultured and co-cultured U2932 cells, (paired T-test).

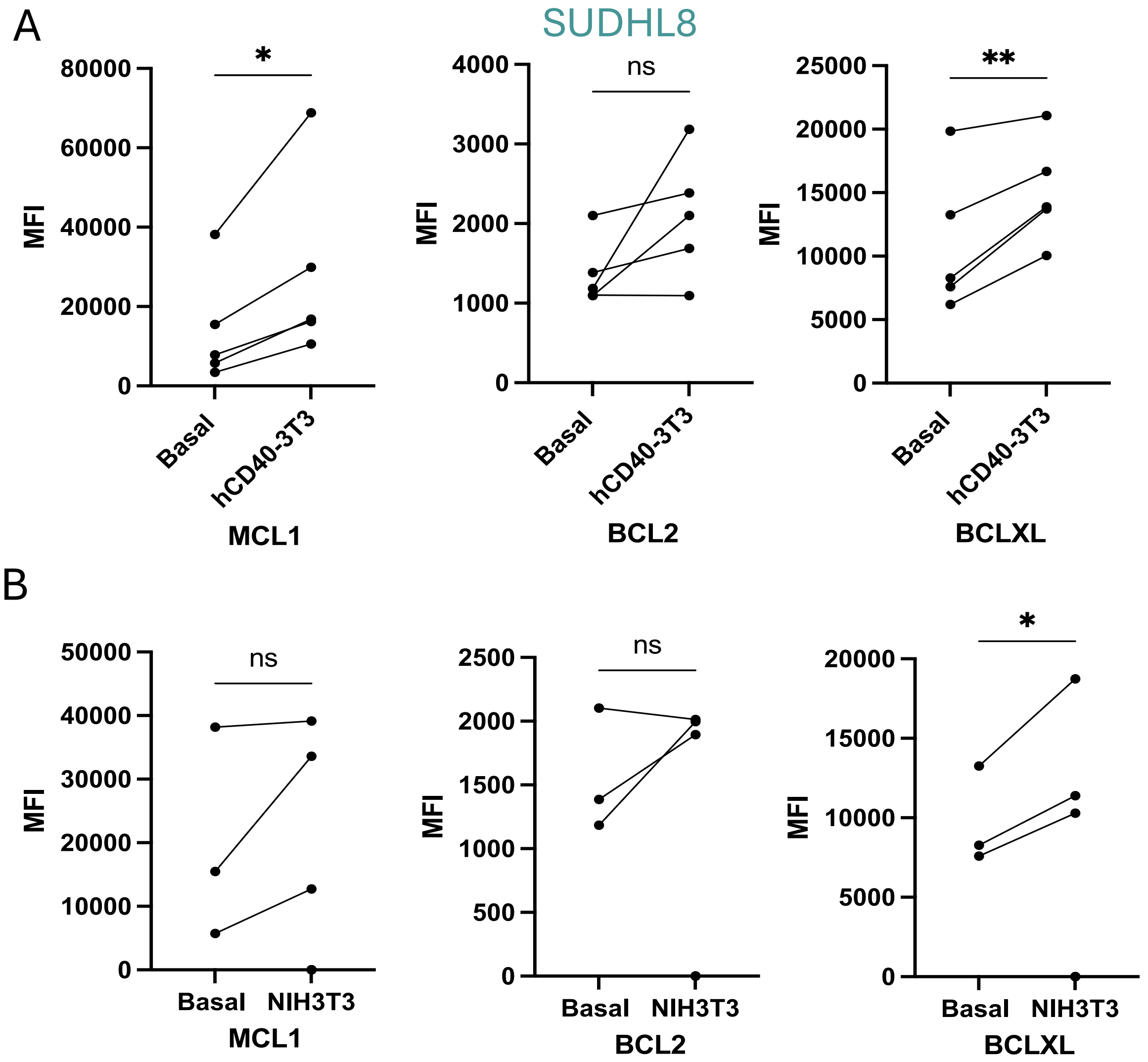


**Supplemental Figure 3.** Higher anti-apoptotic levels are seen in DLBCL cells when are co-cultured with CD40L transfected fibroblasts. (A) SUDHL8 cells were incubated for 24 hours with or without the presence of CD40L transfected fibroblasts. After the stimulation, the DLBCL cells were harvested and stained for Mcl-1, Bcl-2 and Bcl-xL, and then assessed by using flow cytometry. Statistical analysis was performed on five independent experiments' Mean Fluorescence Intensity (MFI) values, between unstimulated and CD40L stimulated SUDHL8 cells, (*$P$<0.05, **$P$<0.01 paired T-test). (B) SUDHL8 cells were incubated for 24 hours with or without the presence of fibroblasts. After the stimulation, the DLBCL cells were harvested and stained for Mcl-1, Bcl-2 and Bcl-xL, and then assessed by using flow cytometry. Statistical analysis was performed on three independent experiments Mean Fluorescence Intensity (MFI) values, between the monocultured and co-cultured SUDHL8 cells, (*$P$<0.05, paired T-test).

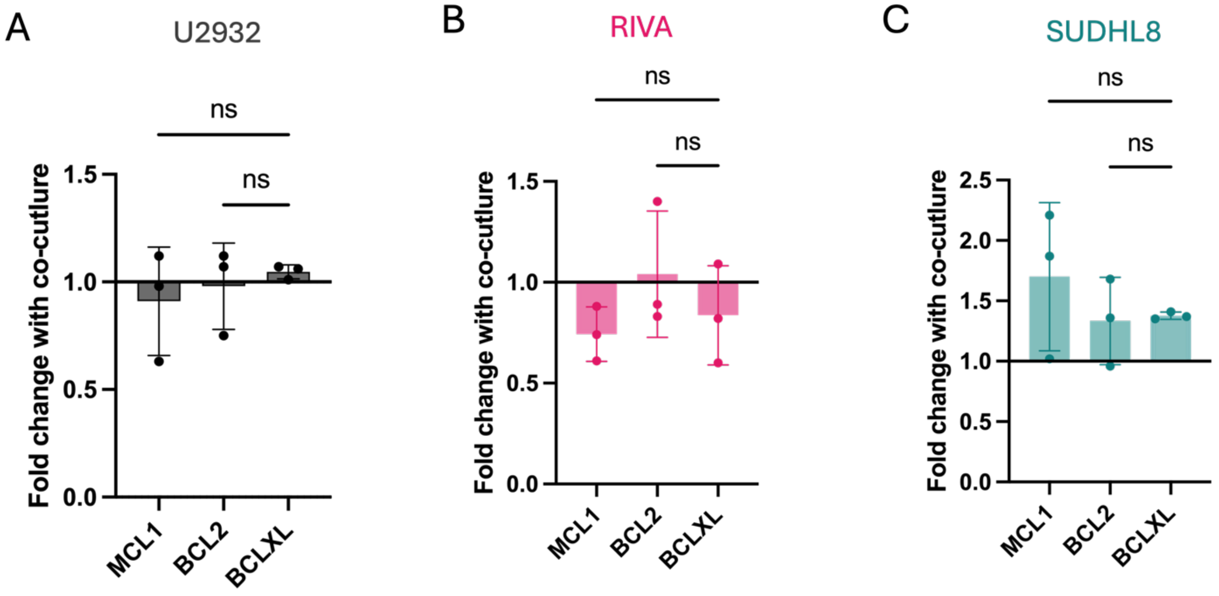


**Supplemental Figure 4.** BCL2-family levels in Diffuse Large B Cell Lymphoma (DLBCL) cell lines, after co-culture with NIH3T3 fibroblasts. (A) MCL1, BCL2, and BCLXL levels, shown as fold changes of 24-hour stimulation with NIH3T3 cells to unstimulated control in U2932 cells with error bars representing the mean ± standard deviation of three independent experiments *(one-way ANOVA with Tukey's comparisons test)*. (B) MCL1, BCL2, and BCLXL levels, shown as fold changes of 24-hour stimulation with NIH3T3 cells to unstimulated control in RIVA cells with error bars representing the mean ± standard deviation of three independent experiments *(one-way ANOVA with Tukey's comparisons test)*. (C) MCL1, BCL2, and BCLXL levels, shown as fold changes of 24-hour stimulation with NIH3T3 cells to unstimulated control in SUDHL8 cells with error bars representing the mean ± standard deviation of three independent experiments *(one-way ANOVA with Tukey's comparisons test)*.

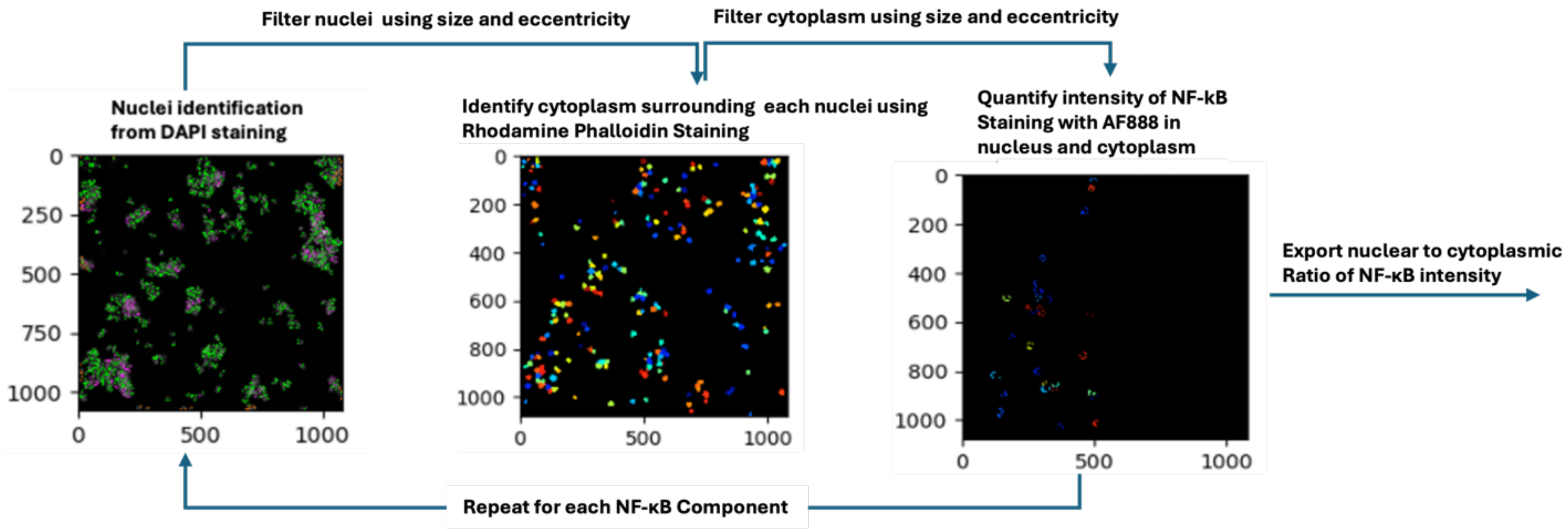


**Supplemental Figure 5.** Automated CellProfiler Pipeline for Nuclear:Cytoplasmic NF-κB analysis. The schematic diagram outlines the analysis performed from the original image. Cells were stained with DAPI to detect the nucleus with the Blue channel, Rhodamin Phalloidin detected with the Red channel to identify the cytoplasm, and AF488 detected from the Green channel to identify NF-κB (RelA, RelB or cRel). Nuclei and cytoplasm were segmented based on size and eccentricity, prior to overlay with AF488 for the quantification of nuclear to cytoplasmic NF-κB ratio for each individual cell with defined nucleus and cytoplasm. Images were acquired on 20x air setting of *OperettaCLS*.

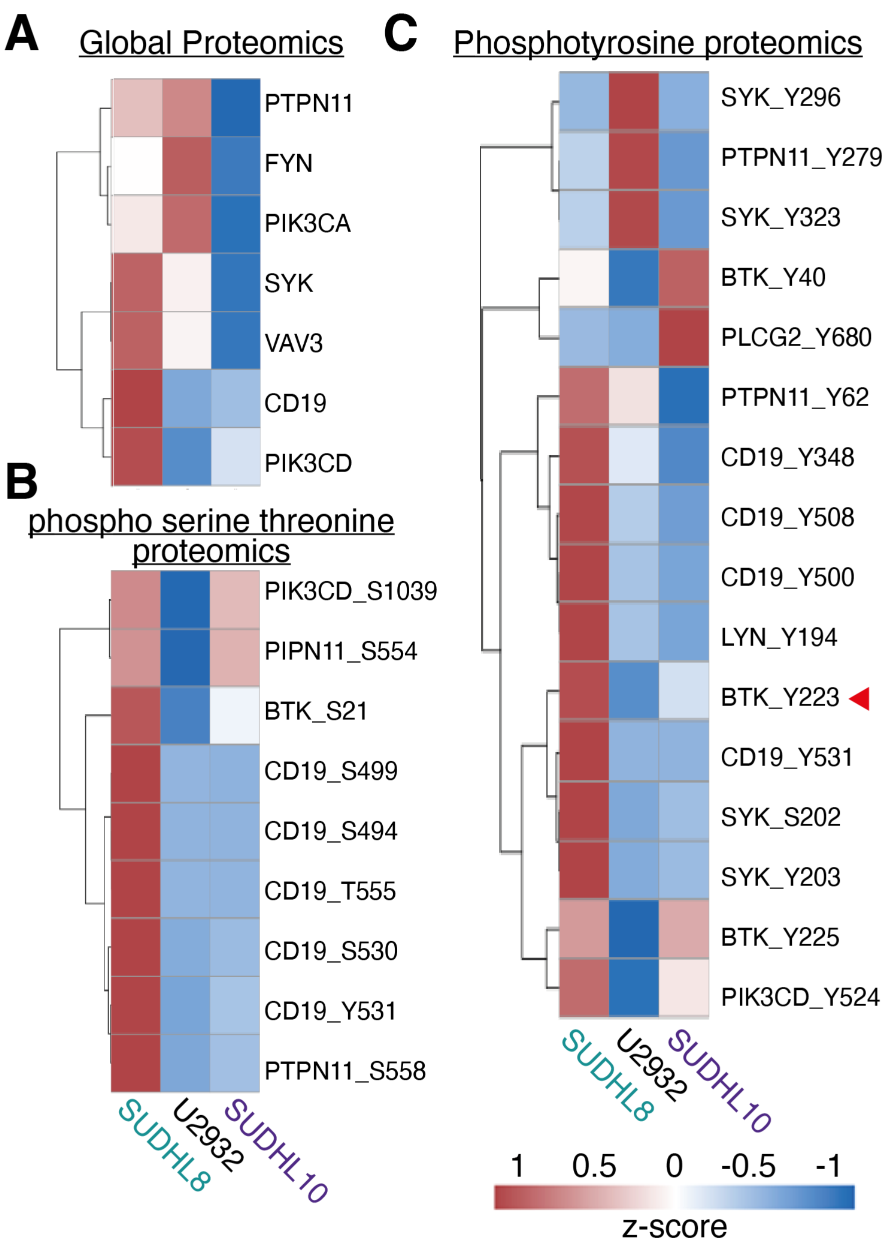


**Supplemental Figure 6.** Heatmap of B Cell Receptor (BCR) Signaling in SUDHL8, U2932, and SUDHL10 Cell Lines. (A) Global proteomics identified 7 BCR-associated proteins with higher expression in SUDHL8 compared to U2932 and SUDHL10. (B) Phospho-serine/threonine proteomics identified 9 BCR-related phosphorylation sites with elevated expression in SUDHL8. (C) Phosphotyrosine proteomics identified 16 BCR-related phosphorylation sites, including BTK_Y223, CD19_Y348, and SYK_Y203, showing higher expression in SUDHL8 compared to the other cell lines.

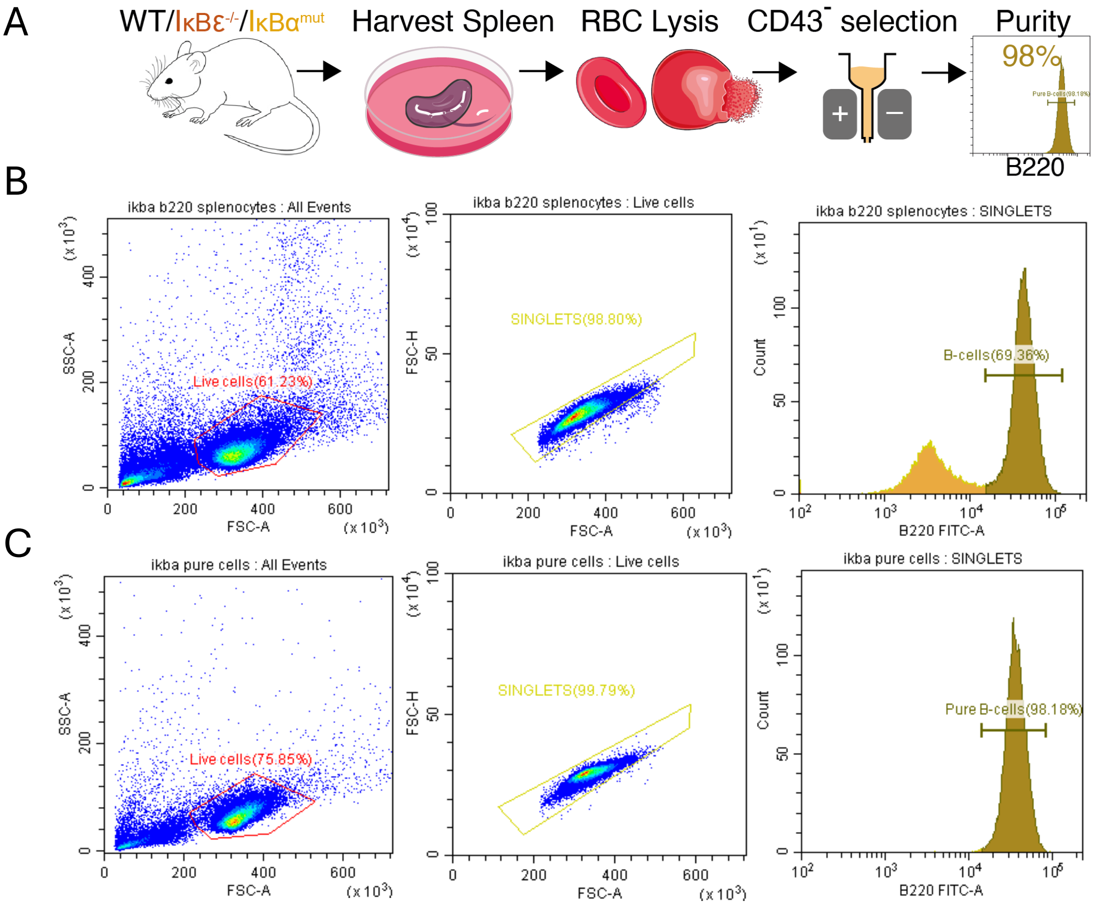


**Supplemental Figure 7.** B cell isolation and purification from primary splenocytes**.** (A) Schematic of B cell isolation and purification from mouse spleens. (B) Flow cytometry gating example on B220⁺ B cells from homogenized mouse splenocytes. (C) B220⁺ gating on mouse B cells, post CD43⁻ magnetic column selection to assess B cell purity percentages.
